# Comprehensive Noise Reduction in Single-Cell Data with the RECODE Platform

**DOI:** 10.1101/2024.04.18.590054

**Authors:** Yusuke Imoto

## Abstract

Single-cell sequencing generates vast amounts of genomic and epigenomic data from thousands of individual cells and can reveal insights into biological principles at the single-cell resolution. However, challenges such as technical noise (dropout) and batch effects hinder obtaining high-resolution structures that are essential for tasks such as the identification of rare cell types and dataset comparison across different cultures. Here, I introduce *integrative RECODE (iRECODE)*, a comprehensive method for noise reduction that is based on the RECODE platform, which targets the technical noise in single-cell RNA-sequencing data using high-dimensional statistics. I show iRECODE effectively mitigates both technical and batch noise with high accuracy and low computational cost. Additionally, the application of RECODE extended to other single-cell sequencing data types including single-cell Hi-C and spatial transcriptomics data and the recent enhancements in RECODE have markedly improved its accuracy and computational efficiency. Thus, the RECODE platform presents a robust solution for mitigating noise in single-cell sequencing, offering promise for advancing our understanding of biological phenomena beyond transcriptomics, encompassing epigenomic and spatial transcriptomic domains.

## 1 Introduction

Single-cell sequencing technologies have driven a paradigm shift in genomics by enabling the resolution of genomic and epigenomic information at an unprecedented single-cell scale. The emergence of expansive single-cell databases such as the Human Cell Atlas [1] and Tabula Sapiens [2] underscores the urgent need for integrating single-cell data analysis across multiple genomic and epigenome datasets.

However, the full potential of these datasets remains unrealized due to technical noise (dropout) and batch effects, which confound data interpretation [3]. Technical noise is a non-biological fluctuation caused by the non-uniformity of detection rates of molecules. It masks true cellular expression variability and complicates the identification of subtle biological signals. Moreover, the high dimensionality of single-cell data introduces the *curse of dimensionality* that obfuscates the true data structure under the effect of accumulated technical noise [4, 5]. Batch effects further exacerbate analytical challenges by introducing nonbiological variability across different datasets, stemming from minute differences in experimental conditions and sequencing platforms. These variations manifest as batch effects that distort comparative analyses and impede the consistency of biological insights across datasets.

Despite the numerous proposed techniques for technical noise reduction (imputation) and batch correction (integration) [6, 7], the simultaneous reduction of both noise types remains a challenge. This shortfall is primarily due to the dependency of conventional batch correction methods on dimensionality reduction, such as PCA, to minimize computational overhead. Such methods require pairwise distance computations between batches, where the computational load scales with dimensionality. In addition, high-dimensional operations can significantly diminish the accuracy of the batch correction owing to the curse of dimensionality. Consequently, simply combining technical noise reduction with batch correction methodologies fails to effectively mitigate both types of noise in single-cell data analyses.

Previously, we developed the RECODE algorithm that modeled technical noise as a general probabilistic distribution, including the negative binomial distribution, and reduced it based on an eigenvalue modification theory in highdimensional statistics [8]. Since then, RECODE has consistently outperformed other representative imputation methods regarding accuracy, speed, and practicability (parameter-free), therefore representing a significant advance in the technical noise reduction of single-cell sequencing data. However, although RECODE was effective in resolving issues associated with technical noise, it neglected the effects of batch noise.

In this study, I establish a comprehensive noise reduction approach that further enhances the RECODE algorithm, making it capable of not only mitigating both technical and batch noise while preserving data dimensions but also processing various types of single-cell sequencing data, including epigenomic and spatial transcriptome datasets. Moreover, by improving its accuracy and computational speed, RECODE is now a versatile platform capable of resolving large-scale integrative single-cell analyses that span multiomics layers and cell types, thereby enhancing our ability to address more complex biological phenomena.

## Results

### Enhancing RECODE for dual noise reduction of single-cell sequencing data

As single-cell sequencing gives rise to two common problems, namely technical noise, which includes dropout events, and batch effects, to simultaneously address these issues I developed iRECODE. This method synergizes the highdimensional statistical approach of RECODE [8] with the established batch correction approach. The original RE-CODE maps gene expression data to an essential space using noise variance-stabilizing normalization (NVSN) and singular value decomposition and then applies principal component variance modification and elimination (Fig. 1a, Methods). As the accuracy and computational efficiency of most batch correction methods decline as the dimensionality increases (Fig. S1), iRECODE was designed to integrate batch correction within this essential space, thereby minimizing the decrease in accuracy and the increase in computational cost by bypassing high-dimensional calculations (Fig. 1b, Methods). This enabled us to implement a simultaneous reduction in technical and batch noise with low computational costs. In addition, iRECODE allows the selection of any batch correction method within its platform.

**Figure 1.**
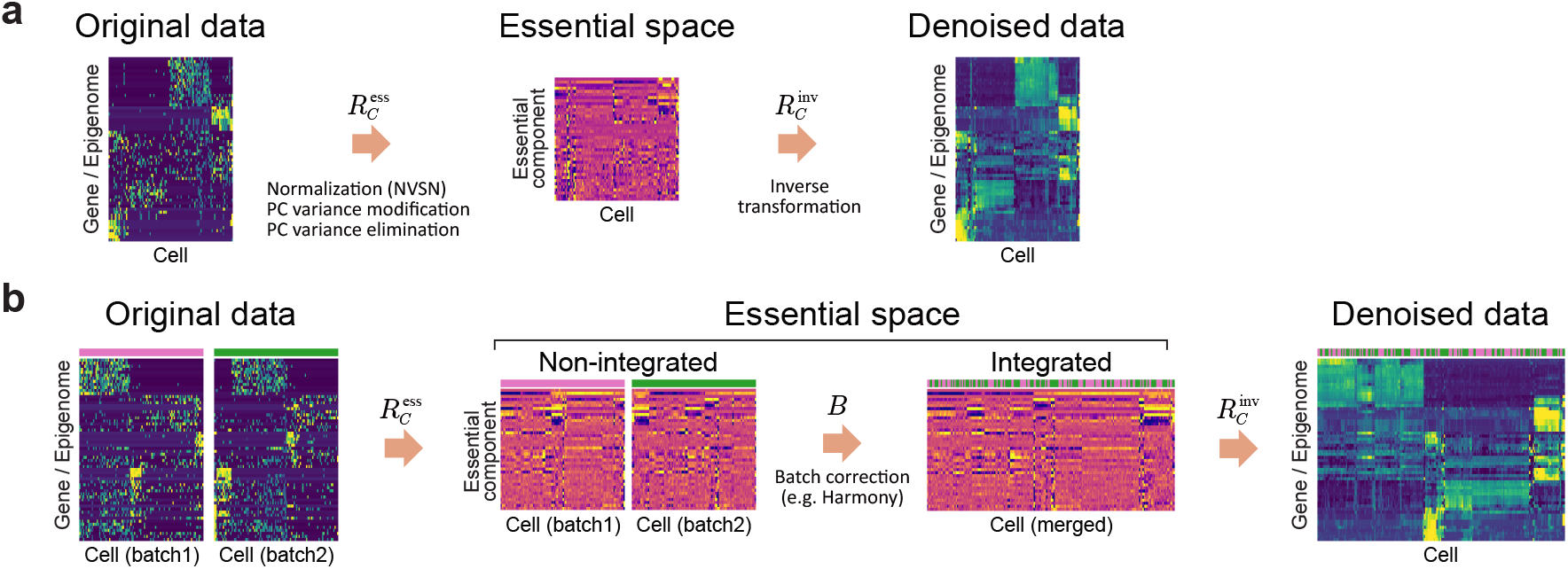
Schematic representation of the RECODE platform for comprehensive noise reduction in single-cell sequencing data. **a**, The RECODE algorithm initiates the noise reduction process by mapping the original single-cell sequencing data into an essential space through noise variance-stabilizing normalization (NVSN). This series of processes denoted as 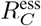, attenuates technical noise via the principal component (PC) variance modification and elimination. The subsequent stage involves an inverse transformation, represented by 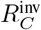, which transposes the denoised data back to the original gene expression space. **b**, Enhancing the RECODE platform, iRECODE incorporates a batch correction step, *B*, within the essential space to address the batch noise. This addition enables us to align multiple batches released from the sparsity and the curse of dimensionality into a singular analytical platform, thus dramatically enhancing the accuracy of downstream data analyses.

Preliminary analysis, using scRNA-seq data comprising three For preliminary analysis, I used scRNA-seq data comprising three datasets and two cell lines [7, 9], to compare the compatibility of three prominent batch correction algorithms –Harmony [10], MNNCorrect [11], and Scanorama [12]– with iRECODE. The result indicated that Harmony performed the best for batch correction (Fig. S2), and I therefore used Harmony as the batch correction in the iRECODE algorithm hereafter. The application of iRECODE successfully mitigated batch effects, as demonstrated by improved cell-type mixing and local inverse Simpson’s index (LISI) metrics, while also releasing the sparsity of the gene expression distribution and reducing dropout rates (Fig. 2a–c). Thus, iRECODE presents a robust approach for the simultaneous reduction of technical and batch noise in single-cell data analyses.

**Figure 2.**
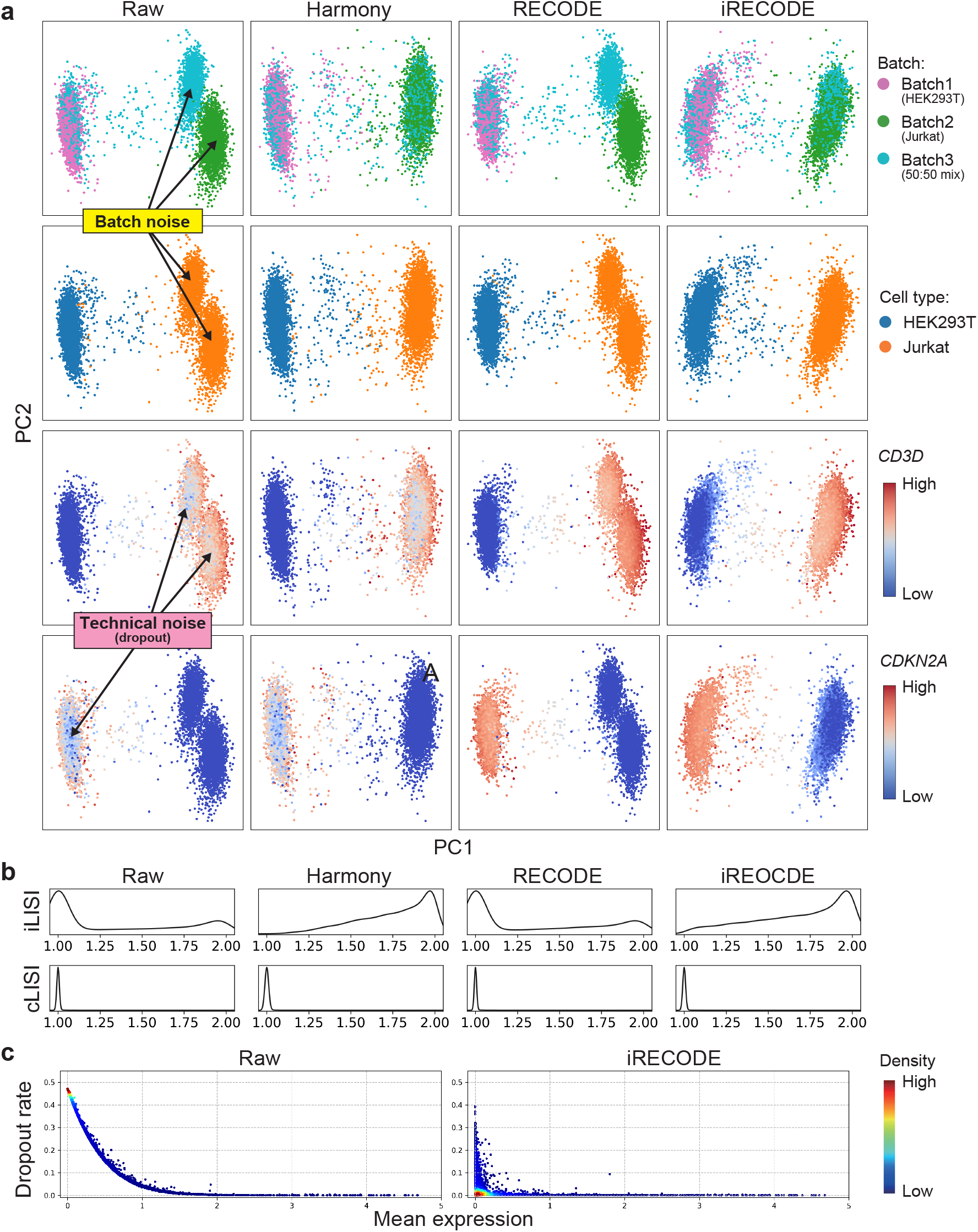
Efficacy of iRECODE in mitigating technical and batch noise in scRNA-seq data. **a**, Principal component analysis (PCA) results of scRNA-seq data comparing three batches and two cell types in various conditions: unprocessed (Raw), processed with Harmony, with RECODE, and with iRECODE. **b**, Assessment of integration quality using distributions of local inverse Simpson’s index for both batch integration (iLISI) and cell type conservation (cLISI). **c**, Dropout rates against mean expression values before and after iRECODE. The raw data show distinct batch effects and technical noise, leading to the non-biological segregation and miss-annotation of clusters by batch and gene expression sparsity. RECODE and Harmony each independently ameliorate a specific aspect of noise, but iRECODE synergistically harmonizes the datasets, resulting in consistent integration across cell types (**b**) and a marked reduction in dropout events (**c**), illustrating the comprehensive capabilities of iRECODE in enhancing data fidelity for downstream analysis.

### Advancing towards authentic single-cell transcriptome analysis with iRECODE

Next, I quantitatively evaluate the performance of iRECODE in comparison to established methods for reducing technical and batch noise using the dataset of the previous section. iRECODE refined the gene expression distributions and accordingly, also addressed dropout and sparsity, mirroring the efficacy of the original RECODE method (Fig. 3a). iRECODE notably modulates the variance among non-housekeeping genes, while consistently diminishing the variance among housekeeping genes, indicating a successful reduction in technical noise (Fig. 3b).

**Figure 3.**
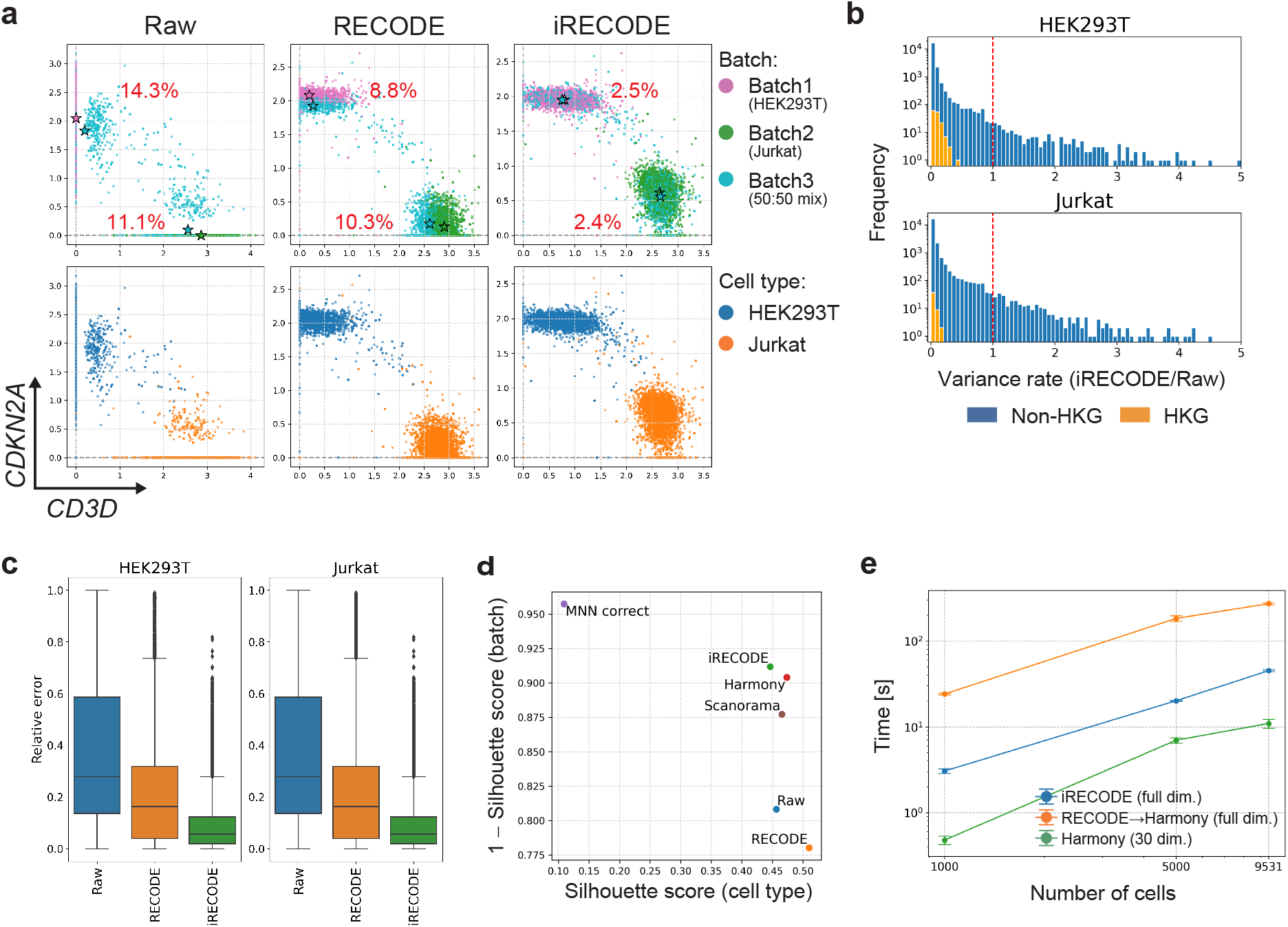
Quantitative assessment and comparative analysis of iRECODE for scRNA-seq data. **a**, Scatter plots presenting gene expression of *CD3D* and *CDKN2A* across different batches and cell types in raw, RECODE, and iRE-CODE processed data. Stars indicate the centroid of gene expression for each group, with percentages reflecting the batch effect-reflected relative errors between centroids per cell type. **b**, Histograms displaying the variance ratio of gene expression between iRECODE integrated and raw data, with blue representing non-housekeeping genes (non-HKG) and orange denoting housekeeping genes (HKG). **c**, Box plots demonstrating the distribution of the relative errors reflecting batch effect for HEK293T and Jurkat cell lines across processing states: raw, RECODE, and iRE-CODE. **d**, Silhouette scores quantifying cell type homogeneity and batch effect mitigation, with the scores indicating improved data integration by iRECODE compared to the raw and prominent batch correction method processed data. **e**, The computational runtime for iRECODE, Harmony, and the combination of original RECODE and Harmony is analyzed across varying cell numbers, illustrating the scalability and efficiency of iRECODE. Input data dimensionality was set as 21,474 genes.

Improvements in batch noise correction were substantial, with the relative errors in the mean expression values significantly decreasing from 11.1–14.3% to a mere 2.4–2.5% (Fig. 3a). On a genomic scale, iRECODE notably enhanced the relative error metrics by over 20% and 10% from those of raw data and traditional RECODE-processed data, respectively (Fig. 3c). Moreover, performance comparisons with established batch correction methods using the silhouette score show that iRECODE is as effective as Harmony, MNN-correct, and Scanorama, demonstrating its accuracy (Fig. 3d).

The precision of iRECODE could also be applied to datasets produced by various scRNA-seq technologies, including Drop-seq, Smart-Seq, and multiple 10x Genomics protocols (Figs. S3 and S4). Despite the greater computational load due to the preservation of data dimensions, iRECODE was approximately ten times more efficient than the combination of technical noise reduction and batch correction methods (Fig. 3e). Togather, this establishes iRECODE as a viable and comprehensive approach for noise reduction in single-cell analysis.

### Application of RECODE to single-cell epigenomics and spatial transcriptomics

The capabilities of RECODE extend beyond scRNA-seq, offering a promising solution for the inherent technical noise present in other data types derived from similar random sampling mechanisms, such as scATAC-seq and scHi-C for epigenomics and spatial transcriptomics. These methodologies rely on random molecular sampling using sequencing, akin to scRNA-seq. This section demonstrates the versatility of RECODE in processing and refining single-cell epigenome and spatial transcriptome data, underscoring its broad applicability and potential to enhance data accuracy across various single-cell sequencing platforms.

### Refining single-cell Hi-C data analysis with RECODE

Single-cell high-throughput chromosome conformation capture (scHi-C) data, presenting a matrix of contact frequencies within chromosomes, offer insights into understanding cell-specific epigenomic architecture distinct from transcriptomic data. However, the sparsity inherent in scHi-C data requires robust noise reduction strategies to enable meaningful cell annotation and significant interaction detection. As noise generated between scHi-C and scRNA-seq datasets are similar, this suggests that RECODE may be effective in reducing noise associated with scHi-C data. To test whether RECODE could discern differential interactions (DIs) that define cell-specific interactions, using sci-Hi-C datasets from five human cell lines at 1M-bp resolution [11], it was applied to the matrix data constructed by vectorizing the upper triangle of scHi-C contact maps. The noise variance-stabilizing normalization (NVSN) distribution, which is an indicator for the applicability of RECODE, revealed that scHi-C data affected by technical noise can be effectively processed using RECODE (Fig. S5a). Indeed, RECODE considerably mitigated data sparsity, aligning the scHi-C-derived topologically associating domains (TADs) with their bulk Hi-C counterparts (Fig. 4a, b). Raw scHi-C data, constrained by variances and coefficients of variation influenced by bin distances and mean values, obscured the heterogeneity of interactions among cells (Fig. S5b). However, RECODE-processed data revealed significant interactions that more accurately reflected this heterogeneity, thereby overcoming the limitations of the raw data. Subsequent PCA and UMAP analyses of the processed data confirmed distinct cell type segregation (Fig. 4c, d). Remarkably, the performance of RECODE was on par with scHi-C data-specialized preprocessing methods [13, 14], despite using the same noise reduction platform as the single-cell transcriptome data.

**Figure 4.**
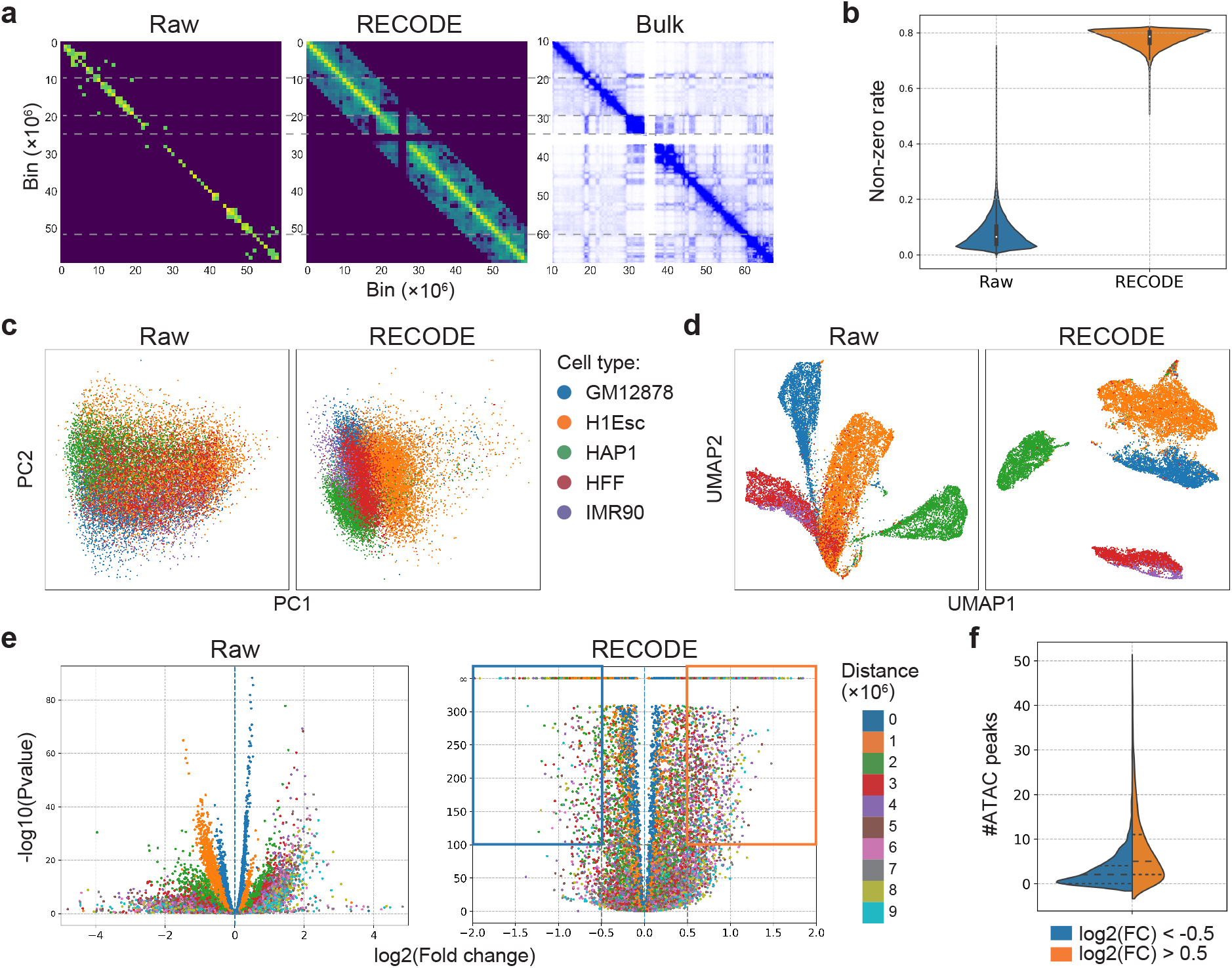
Evaluation of RECODE’s impact on scHi-C data interpretation. **a**, Comparison of contact maps at 1M-bp resolution for raw, RECODE-processed scHi-C data, and bulk Hi-C in GM12878 cells, delineating chromosomal interactions at chromosome 19. **b**, Violin plots illustrating the non-zero interaction rate for raw versus RECODE-processed scHi-C data, highlighting RECODE’s ability to alleviate data sparsity. **c, d**, Dimensionality reduction visualizations, with PCA and UMAP, for raw and RECODE-processed scHi-C data, colored by cell type, indicating enhanced data segregation post-RECODE processing. **e**, Volcano plots contrasting differential interactions (DIs) within GM12878 cells against other cell types, with color coding based on inter-bin distances. These plots emphasize the distribution of significant interactions, with the right section (log2(FC) *>* 0) spotlighting GM12878-specific DIs. Interaction specificity for GM12878 cells is marked by the highlighted orange and blue box regions. **f**, Distribution of the count of ATAC peaks for GM12878 cells, categorized based on bins within boxes in **e**, demonstrating the disparity in chromatin accessibility associated with DIs.

In the downstream DI analysis, while raw scHi-C data struggled to detect significant DIs owing to the drop in detection rate corresponding to the distance between bins, RECODE precisely identified cell type-specific DIs that overlapped with ATAC-seq-derived open chromatin regions (Fig. 4e, f). Notably, RECODE showed chromosome 19 had significant cell type-specific interactions(Fig. S5c, d). Thus, RECODE effectively denoises single-cell epigenome sequencing data under the same principles used for single-cell transcriptome data, thereby revealing intricate cell-specific chromosomal and epigenomic landscapes.

### Refining spatial transcriptome analysis with RECODE

Spatial transcriptome analysis significantly enhances our understanding of cellular heterogeneity and interactions within native tissue environments and provides critical insights into the complex relationships of biological systems. Similar to other sequencing technologies, it faces the challenge of technical noise, which can mask biological signals and obstruct key discoveries. Because the gene expression part of the spatial transcriptome data is generated through mechanisms similar to those of other sequencing technologies, the application of RECODE, a method for denoising the technical noise generated in the sequencing process, can effectively resolve this issue.

I applied RECODE to gene expression data of spatial transcriptome datasets derived from the Stereo-seq and 10x Visium platforms. Based on the NVSN distributions, gene expression data from the spatial transcriptome datasets contained RECODE-modeled technical noise (Fig. S6). Although Stereo-seq achieves near-single-cell resolution, raw data sparsity remains an issue (Fig. 5a, b). However, the application of RECODE refines these distributions, revealing distinct spatial gene expression patterns. This resolution enabled the identification of spatial functions for genes with low expression levels, such as *Foxc2, Gata4*, and *Grhl2* (Fig. 5b). Similar sparsity challenges in 10x Visium data were resolved by RECODE, clarifying gene expression within spatial tissues, regardless of the cell type or species (Fig. 5c–d). For instance, RECODE aids in delineating spatial patterns of cancer progression by highlighting cancer-specific gene expression (Fig. 5d). Hence, the application of RECODE successfully revealed intricate spatial gene expression patterns, shedding light on the potential function of those spatially specific genes.

**Figure 5.**
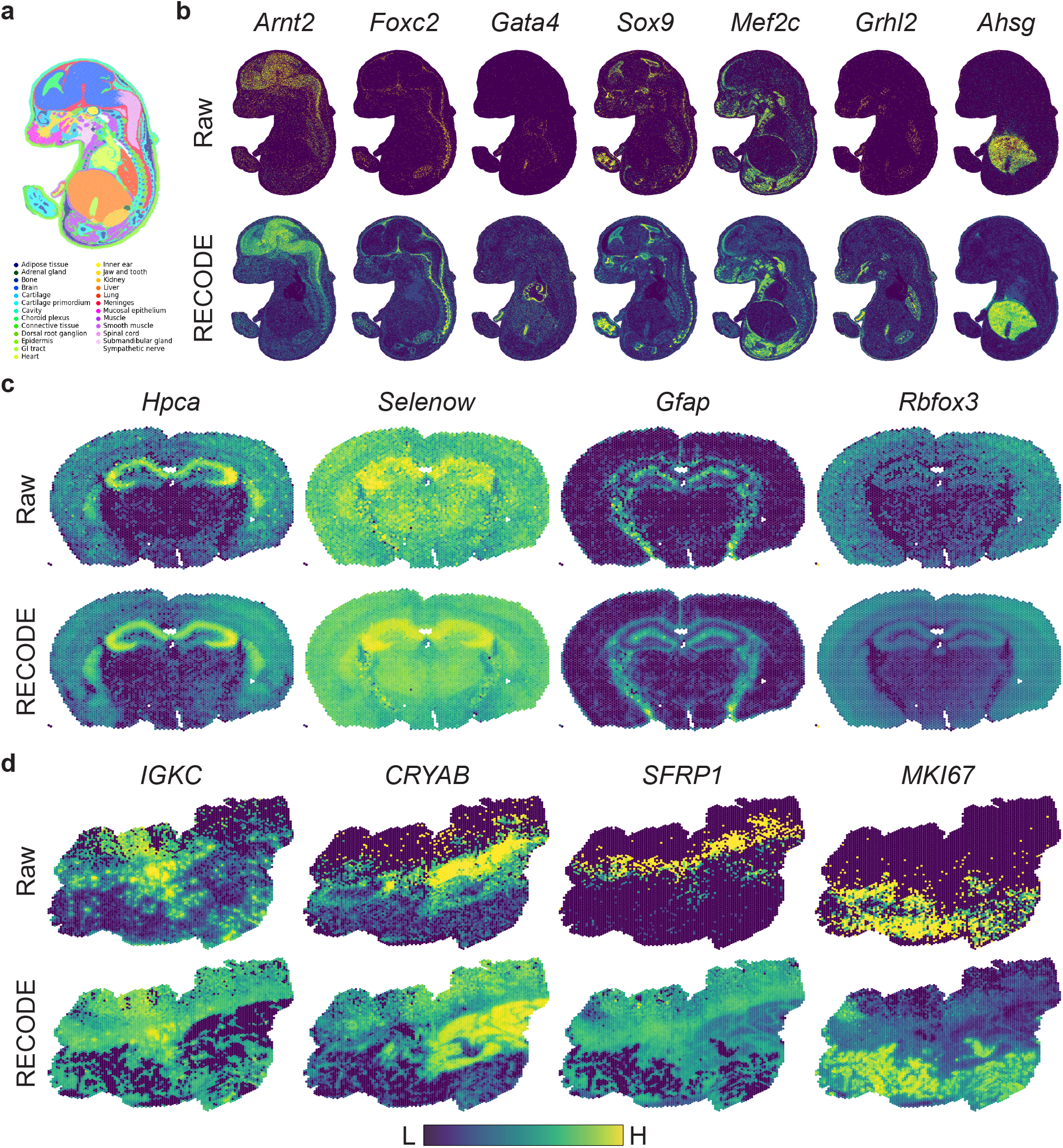
Illustration of RECODE’s denoising efficacy on spatial transcriptomic data. **a**, Annotated cell type spatial distribution within a mouse embryo at embryonic day 16.5, delineating the complex tissue structure. **b**, Side-by-side visualization of spatial gene expression patterns for selected cell-type-specific markers in the mouse embryo, contrasting raw data with RECODE-processed data. **c** and **d**, Comparative spatial gene expression profiles highlighting cell identity in mouse brain (**c**) and human colon cancer (**d**) tissue sections. The upper panels correspond to raw data visualizations, whereas the lower panels showcase the enhanced clarity following RECODE processing. These comparative visualizations underscore the capacity of RECODE to enhance data interpretability by reducing background noise and amplifying biologically relevant signals across distinct tissue types.

### Enhancing accuracy and reducing computational costs of RECODE

Recent advances in high-dimensional statistical theory have substantially bolstered the noise reduction capabilities of RECODE. By incorporating Yata Aoshima’s theoretical eigenvalue modification [15, 16], the original RECODE addresses the curse of dimensionality [8]. They also introduced a novel approach to modify eigenvectors using sparse PCA theory [17], which I integrated into the updated version of RECODE. This eigenvector modification, occurring simultaneously with the eigenvalue modification within the essential space of RECODE, further mitigates technical noise that remains in the essential space of RECODE (Methods). This update results in the preservation of variation in marker genes and a reduction in housekeeping genes, indicating an enhancement in the detection of key differentially expressed genes (Fig. 6a–b).

**Figure 6.**
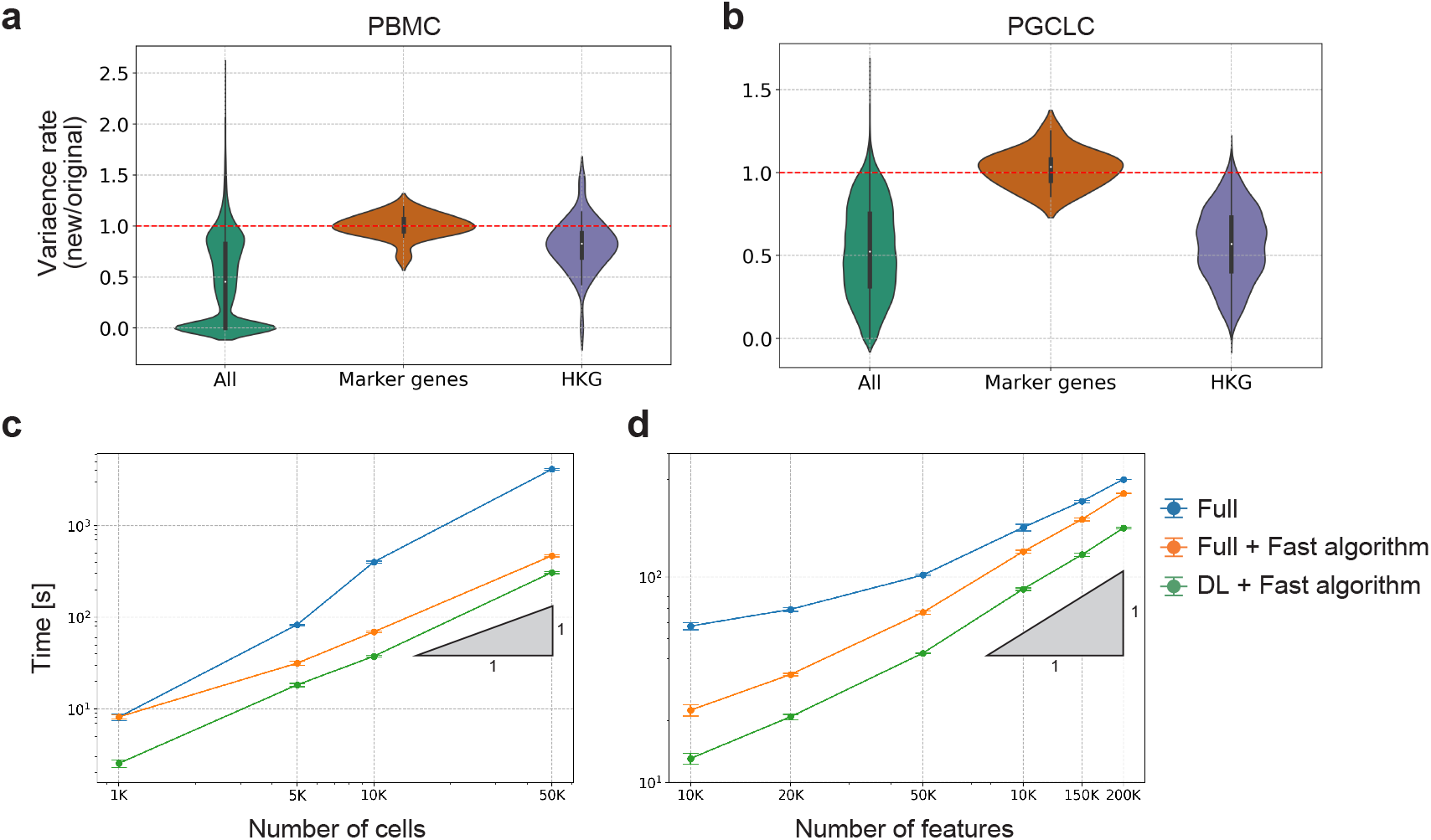
Assessment of RECODE’s enhanced algorithmic accuracy and computational performance. **a, (b)**, Violin plots of variance ratios for PBMC and PGCLC datasets, respectively, comparing gene expression variability across all genes, marker genes, and housekeeping genes (HKG) between the original and the improved RECODE algorithm with eigenvector modification. The dashed red line indicates equal variance between the two versions. **c, d**, Computational time required by the RECODE algorithm relative to the number of cells and features analyzed. Performance metrics compare the naive algorithm using all cells (full), an established fast algorithm (Methods), and a downsampling learning (DL) strategy combined with the fast algorithm. Standard deviations are denoted by error bars, while the triangles represent linear scalability in processing time. These visualizations quantify the advancements in RECODE’s algorithmic accuracy for biological marker gene detection and its computational efficiency, pivotal for large-scale single-cell data analysis.

To further improve the practicality of RECODE, I also incorporated a downsampling learning (DL) approach. This allows RECODE to learn denoising functions from a randomly selected subset of single-cell data, thereby preserving the need for full dataset utilization (Methods). By applying this fast RECODE algorithm, computational efficiency was greatly improved without sacrificing accuracy. With a modest 20% of the data employed in the learning phase, the DL algorithm markedly accelerated the operational speed of RECODE, outpacing other previous computations (Fig. 6c–d). Moreover, this algorithm showed near-first-order scalability concerning both the cellular and feature dimensions (Fig. 6c–d), thereby ensuring efficiency in the face of emerging big datasets.

## Discussion

In this study, I developed an integrative preprocessing methodology for single-cell sequencing data, based on the advancement of our noise reduction technique, RECODE. Grounded on the principles of high-dimensional statistics, RECODE addresses the pervasive issue of technical noise commonly associated with dropout events in single-cell sequencing data. The evolved methodology, which I have called iRECODE, expands this capability to encompass batch noise attenuation. This is achieved by integrating batch correction directly within the algorithm’s essential low-dimensional space, giving rise to a method that not only improves the precision of noise reduction, but also substantially decreases computational demand. Additionally, Harmony was found to be the most compatible for batch correction with iRECODE, possibly due to it not assuming a gene expression space, therefore allowing for batch correction in RECODE’s essential space.

I also showed the utility of RECODE goes beyond transcriptomics, encompassing both single-cell epigenomics and spatial transcriptomics, by distinguishing cell types using chromosomal architecture and resolving spatial patterns of low-expression genes, respectively. Furthermore, enhancements in the accuracy and processing speed of RECODE have further enabled it to handle the influx of large-scale datasets, ensuring faster and more precise noise reduction. These advances significantly broaden the applications for RECODE, establishing it as a fundamental tool that will become the foundation for single-cell data analysis across extensive datasets.

Looking back at the theory of RECODE, it models the data generation of single-cell sequencing as random sampling, removing noise while preserving the true gene expression distribution, thereby ensuring no loss of genuine biological signals. This approach avoids the *cyclicity* issue commonly observed in traditional imputation methods, in which improvements in certain aspects inadvertently lead to detrimental effects on others [3]. Additionally, RECODE is applicable to a wide range of sequencing platforms because of noise modeling inherent in the sequencing process, without limiting specific molecules, cell types, or data generation platforms. Furthermore, as RECODE retains the original data structure, including the matrix size and expression scale, the data processed by RECODE can be seamlessly integrated with conventional normalization and downstream analyses. Therefore, RECODE will likely emerge as an essential preprocessing step in analyzing all single-cell data in the future.

Nevertheless, RECODE also has its current limitations. In scenarios such as early cell differentiation or cancer cells with relatively few mutations, where genetic information is overwhelmingly similar, traditional clustering may lead to errors even when non-biological noise is completely removed. Therefore, in post-noise reduction, the selection of biologically variable genes for classification becomes pivotal, potentially requiring supplementary methodologies such as ScType [18].

Furthermore, distinguishing batch noise from slight biological variations remains a critical challenge. However, the flexibility in choosing batch-correction methods in iRECODE suggests that as research in this area progresses, so will the capabilities of iRECODE.

Despite its current capabilities, further refinement of applications for RECODE will important future goal. Some datasets classified as weakly applicable indicate the presence of nonrandom sampling noise (Table S1). One potential solution is the mathematical enhancement of the noise variance model, which can further improve the utility of RE-CODE. Additionally, the application of RECODE to diverse genomic and epigenomic datasets is expected to reveal biological insights that were previously concealed by noise.

In sum, RECODE is a comprehensive and versatile platform for noise reduction in single-cell sequencing, with the potential to go beyond its current transcriptomic applications in epigenomics and spatial transcriptomics, revolutionizing our understanding of complex biological systems.

## Acknowledgements

I extend my sincere appreciation to Drs. Cantas Alev, Yasuaki Hiraoka, Masahiro Nagano, Tomonori Nakamura, Taro Tsujimura, and Kazuyoshi Yata for their invaluable discussions and insights, which have greatly contributed to the advancement of this work. I would also like to thank Dr. Spyros Goulas for critical comments and constructive suggestions on my manuscript. This research was supported by the World Premier International Research Center Initiative (WPI), Japan, the JST PRESTO program (grant number JPMJPR2021), and the JST FOREST program (grant number JPMJFR222X), which provided funding and an inspirational research environment.

## Author Contributions

The sole author, Yusuke Imoto, conceived the study, designed the methodology, conducted the research and analysis, and wrote the manuscript.

## Methods

### RECODE

Here, we provide an overview of the RECODE algorithm [8]. We first introduced noise variance-stabilizing normalization (NVSN), which is the first step in the RECODE algorithm. Let ℕ, ℕ_0_, and ℝ be sets of natural numbers, non-negative integers, and real values, respectively. Let 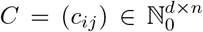 be the count data matrix, such as the raw single-cell genome and epigenome sequencing data, which are the inputs of RECODE, where *d* and *n* are the dimensions ( number of features) and number of cells, respectively. We define 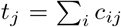 as the total count of cells *j* and remove cells with *t*_*j*_ = 0 from the input data in advance. Then, we define NVSN *F*_*C*_ : ℝ^*d×n*^ *→* ℝ^*d×n*^ as

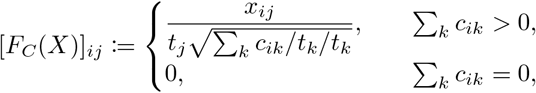

where *X* = (*x*_*ij*_) ∈ ℝ^*d×n*^. We introduced *F*_*C*_(*C*) the NVSN matrix.

Next, the transformation of the RECODE algorithm is introduced. For matrix *X* = (*x*_*ij*_) ∈ ℝ^*d×n*^, let *S*_*X*_ := (*X* − *XP*)(*X* − *XP*)^T^*/*(*n* − 1) ∈ ℝ^*d×d*^ be the covariance matrix of *X*, where *P* is an *n × n* matrix whose elements are all 1*/n* (*XP* shows the row-wise mean vector). Further, let *U*_*X*_ = (*u*_*X*,1_, …, *u*_*X,d*_) ∈ ℝ^*d×d*^ and Λ_*X*_ = diag(*λ*_*X*,1_, …, *λ*_*X,d*_) ∈ ℝ^*d×d*^ (*λ*_*X*,1_ ≥ … ≥ *λ*_*X,d*_) be the components of the singular value decomposition of *S*_*X*_, i.e., 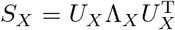. Here, (*λ*_*X,i*_, *u*_*X,i*_) denotes the pair of *i*th eigenvalue and eigenvector of the covariance matrix *S*_*X*_. Furthermore, *u*_*X,i*_ and *λ*_*X,i*_ correspond to the *i*th axis and variance of PCA, respectively. Yata and Aoshima proposed a modification of the PC variance as

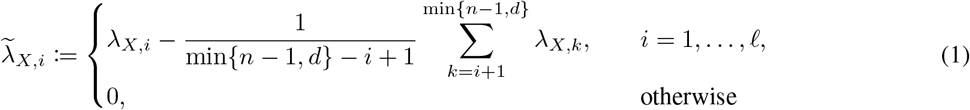

and proved an improvement in its convergence in a high-dimensional statistical setting [16]. Here, 𝓁 ≤ min*{n* − 1, *d}* is a parameter called the essential dimension of the matrix *X*. We propose the optimal value 𝓁* of the essential dimension for the normalized matrix *F*_*C*_(*C*) as:

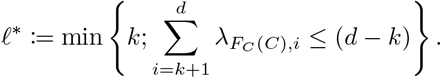

Then, we define the RECODE transformation *R*_*C*_: ℝ^*d×n*^ *→* ℝ^*d×n*^ for the count data matrix *C* as:

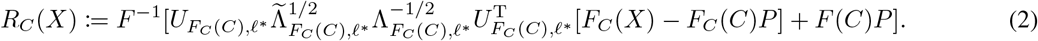

Here, 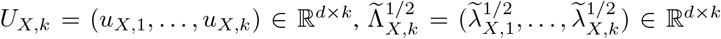, and 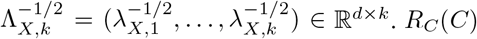 is the RECODE output matrix.

### Applicability

RECODE determines the suitability of noise reduction efforts through a robust analysis of gene variance within the NVSN matrix, stratifying datasets into three distinct categories: strongly applicable, weakly applicable, and inapplicable [8]. Strong applicability indicates congruence with the RECODE noise model, which is conducive to substantial technical noise mitigation. Weak applicability, while still within the RECODE platform, suggests the presence of additional non-modeled noise. However, inapplicability indicates a fundamental discrepancy with the model, in which the noise deviates from that accounted for by random sampling. This systematic categorization informs the tailored application of RECODE across diverse single-cell sequencing datasets with detailed Table S1.

### Fast algorithm

The fast RECODE algorithm was implemented using the algebraic characteristics between the matrices and eigenvalues [8]. Calculating the average of the subsequent eigenvalues in the traditional PC variance modification requires a full PCA computation. We derived an equivalent algorithm that utilized only a subset of eigenvalues by exploiting the relationship between the trace of the matrix and the eigenvalue. This innovation permits the use of truncated PCA instead of a full PCA, thereby substantially enhancing the computational efficiency of RECODE. Moreover, the combination of a fast algorithm and downsampling learning introduced in the subsequent section represents a significant leap forward in the speed of the algorithm, streamlining the noise reduction process for large-scale single-cell datasets.

### iRECODE

iRECODE unifies multiple datasets in the essential space of RECODE. Using transformations 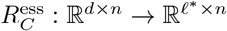 and 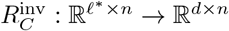, defined as

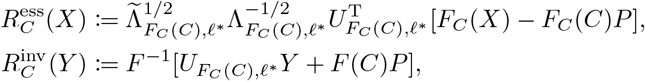

RECODE transformation *R*_*C*_ is represented as 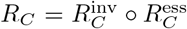. We call 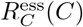 the essential matrix of RECODE. Batch correction in the essential space reduces the computational cost because the essential dimension 𝓁^*^ is usually 10 ∼ 1,000. In summary, using batch correction 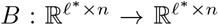, iRECODE transformation *iR*_*C*_ is formulated as

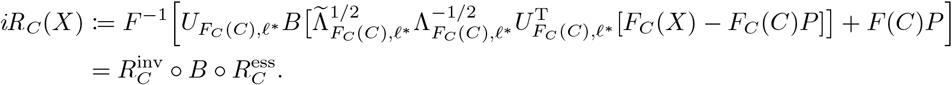

*iR*_*C*_(*C*) is the output matrix of the iRECODE.

### Eigenvector modification for accuracy improvement

RECODE employs an eigenvalue modification of the covariance matrix as a pivotal step in the PC variance adjustment, a method based on the study by Yata and Aoshima [15, 16]. A recent methodological advancement by Yata and Aoshima introduced an eigenvector modification technique [17] that leverages the principles of sparse PCA. This technique refines the eigenvectors by automatically optimizing the parameters through an eigenvalue modification framework. We integrated this cutting-edge approach to increase the accuracy of the RECODE algorithm.

We formulated an eigenvector modification and combined it with the RECODE algorithm. Recall that *λ*_*X,i*_ and *u*_*X,i*_ are *i*th eigenvalue and eigenvector of covariance matrix *X* ∈ ℝ^*d×n*^, respectively. Furthermore, *λ*_*X,i*_ is the modified eigenvalue defined in Eq. (1). For each eigenvector *u*_*X,i*_ = (*u*_*X,i*1_, …, *u*_*X,id*_), we define a symmetric group *σ* : *{*1, …, *d} → {*1, …, *d}* such that |*u*_*X,iσ*(1)_| *≥ · · · ≥* |*u*_*X,iσ*(*d*)_|. We set integer *k*_*i*_ such that

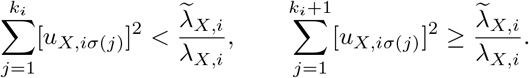

We then define the modified eigenvector 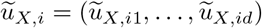 as

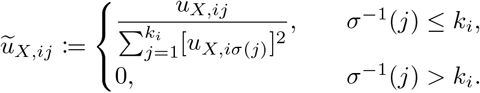

This transformation eliminated the components of the eigenvector that were close to zero based on the rate of modification of the eigenvalues. Using modified eigenvector matrix 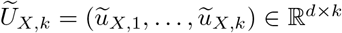, we define accuracyimproved RECODE transformation 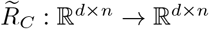 as

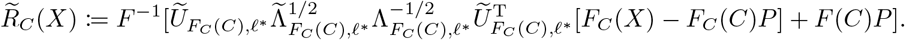

In addition, we introduced an accuracy-improved iRECODE *iR*_*C*_ by replacing 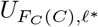 with 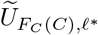. Accuracy improvement was executed using the `version=2` option on the GitHub code.

### Downsampling learning (DL) for acceleration

RECODE operates as an unsupervised learning algorithm that learns a transformation from the input count matrix *C* and applies it to the input data (refer to Eq. (2)). The RECODE algorithm incurs most of the computational costs during the learning process. To enhance efficiency, we implemented a downsampling strategy to expedite learning.

Consider *n*^*^ (where *n*^*^ *< n*) as the sample size after post-downsampling, and 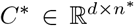 as the corresponding downsampled count data. The downsampling learning algorithm for RECODE, denoted as 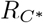, learns transformations using *C*^*^ and subsequently applies these transformations to the original input data *C*, producing denoised data 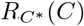. Our GitHub repository for RECODE defaults to a 20% downsampling rate for datasets exceeding 20,000 samples.

### Integration analysis

In our integration analysis, I used benchmark datasets 4–7 from Tran et al. [7]. These included dataset 4: human pancreas analyzed via multiple protocols (Fig. S2c and Fig. S3c), dataset 5: human pancreas using 3’ and 5’ 10x Chromium protocols (Fig. S2b and Fig. S3b), dataset 6: cell line comparison (HEK293T and Jurkat) via Drop-seq (Figs. 2 and 3), and dataset 7: mouse retina by Drop-seq (Fig. S2a and Fig. S3a). iRECODE was applied to raw scRNA-seq data.

The dropout rate ( Fig. 2c) was quantified as the percentage of zero counts per gene. The relative error RE(*i*) for the *i*th gene between batches b1 and b2 (shown in Fig. 3a and c) is defined as

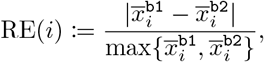

Where 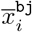 is the mean expression value of the *i*th gene in the *j*th batch. The relative errors were assessed for each cell type. Housekeeping genes (analyzed in Fig. 3b) were identified using the HRT Atlas v1.0 [19]. To evaluate computational efficiency (Fig. 3e), the runtime was measured ten times for randomly selected cells (*n* = 1,000, 5,000, 9,531 (full)). The quantitative integration scores, including LISI and silhouette scores, are as follows:

### LISI

The local inverse Simpson’s index (LISI) quantitatively assesses the efficacy of batch correction by measuring the local diversity around each cell [10]. It is predicated on the variance in annotations among a cell’s nearest neighbors. Integration LISI (iLISI) utilizes batch labels with distributions approaching the number of batches per cell type, indicating thorough mixing. Conversely, cell-type LISI (cLISI) is based on cell-type annotations, where distributions near one suggest the successful segregation of cell types.

### Silhouette score

The silhouette score was used to calculate the goodness of the clustering technique. Its value ranges from −1 to 1, where a high value indicates that the object is well-matched to its own cluster and poorly matched to neighboring clusters. If most objects have high values, then the clustering configuration is appropriate. If many points have low or negative values, the clustering configuration may contain too many or too few clusters. The silhouette score is particularly useful when the ground truth of the data is unknown, and it provides a succinct graphical representation of how well each object lies within its cluster.

### scHi-C data analysis

We utilized single-cell combinatorial-indexed Hi-C (sci-Hi-C) data at 1M-bp resolution from five human cell lines: GM12878, H1Esc, HFF, IMR90, and HAP1, as reported in [13]. The observed contacts were limited to those within 10M-bp inter-bin distances. A matrix was constructed from the cell’s scHi-C contact map by flattening the upper triangular portion. Matrix data were transformed using the RECODE algorithm.

Following RECODE processing, both raw and processed data were subjected to log normalization, setting the stage for downstream analyses. In addition, a bulk Hi-C contact map, as shown in Fig. 4a, was generated using Juicebox [20]. In Fig. 4f, I used bulk ATAC-seq data (ENCSR095QNB) of GM12878 cells downloaded from ENCODE3 data portal.

### Spatial transcriptome analysis

We employed Stereo-seq data from mouse embryos at E16.5 [21] (`E16.5 E1S1.MOSTA.h5ad`) and demonstration data from 10x Visium for mouse brain and human colon cancer. RECODE was applied to the gene expression count matrices excluding the spatial coordinates. Similar to the approach used for scRNA-seq data, standard log-normalization was performed, followed by mapping of the processed gene expression values onto spatial coordinates. The efficacy of RECODE for Stereo-seq and 10x Visium data was determined to be weak and strong, respectively (Fig. S6).

### Verifications of accuracy and computational costs

For accuracy verification, I employed scRNA-seq data of peripheral blood mononuclear cells (PBMC) derived from 10x Genomics demonstration datasets for Fig. 6a and human primordial germ cell-like cell (hPGCLC) induction system data on day 2 [22] for Fig. 6b. Marker genes for PBMCs included *CD14, CD79A, CD8A, CD8B, CST3, FCER1A, FCGR3A, GNLY, IL7R, LGALS3, LYZ, MS4A1, MS4A7, NKG7, S100A8, KLRB1*, and for hPGCLC system *GATA2, GATA3, GATA4, HAND1, MIXL1, NANOG, NANOS3, POU5F1, PRDM1, SOX17, TFAP2A, TFAP2C*, selected based on previous research [22, 23, 24]. Housekeeping genes were identified using HRT Atlas v1.0 [19].

For computational efficiency verification, I employed fresh scRNA-seq data of 68k PBMCs [9] for Fig. 6c and scATAC-seq data of a 5k mix of human GM12878 and mouse A20 cells for Fig. 6d generated using the 10x Chromium platform. Runtime measurements were conducted ten times for randomly selected cells or ATAC regions.

## Data availability

The integration benchmark datasets (datasets 4, 5, 6, and 7) were obtained from Tran et al. [7]. The sci-Hi-C data presented by Kim et al. [13] are available at here. Chen et al. [21] used mouse embryo STEREO-seq data available at MOSTA database. The 10x Visium datasets for mouse brain and human colon cancer are available via the 10x Genomics website, with direct links for mouse brain and human colon cancer. The scRNA-seq data of PBMCs are available from 10x Genomics demo data, and the hPGCLC induction system data are hosted in GEO under accession GSE140021. Zheng et al. [9] scRNA-seq data of 68k PBMCs from the Short Read Archive under accession SRP073767. The scATAC-seq data of a 5k mixture of freshly frozen human GM12878 and mouse A20 cells were obtained from 10x Genomics demo data.

## Code availability

The Python and R implementations of the RECODE algorithm are publicly accessible at GitHub repository. The repository includes the iRECODE feature as a part of the Python codebase.

**Figure S1:**
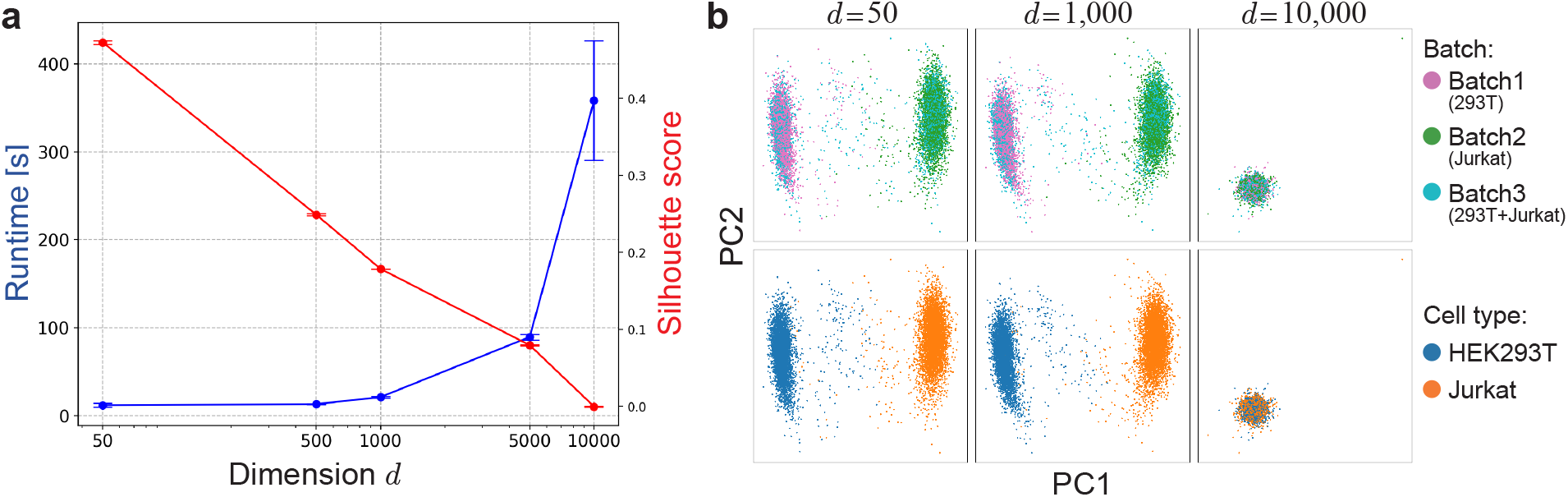
Impact of dimensionality on the performance of batch correction in single-cell data analysis. **a**, A dual-axis plot demonstrating the performance of the batch correction method, Harmony, on scRNA-seq data from three batches containing HEK293T and Jurkat cell lines. The computational runtime, indicated by the blue line and plotted against the left vertical axis, shows an exponential increase with the dimensionality of the data. Conversely, the silhouette score for cell type, indicating the accuracy of batch correction, depicted by the red line and plotted against the right vertical axis, decreases as dimensionality increases, indicating that the batch correction method is also affected by the curse of dimensionality. The error bars represent standard deviations from 10 independent computations. **b**, PCA projections illustrating the separation of cell types and batches at different data dimensionality. The plots reveal that higher dimensionality can degrade the clarity of cell type discrimination within batch-corrected data. These findings emphasize the necessity of dimensionality reduction as a preprocessing of batch correction, highlighting the difficulty in combining batch correction and technical noise reduction methods.

**Figure S2:**
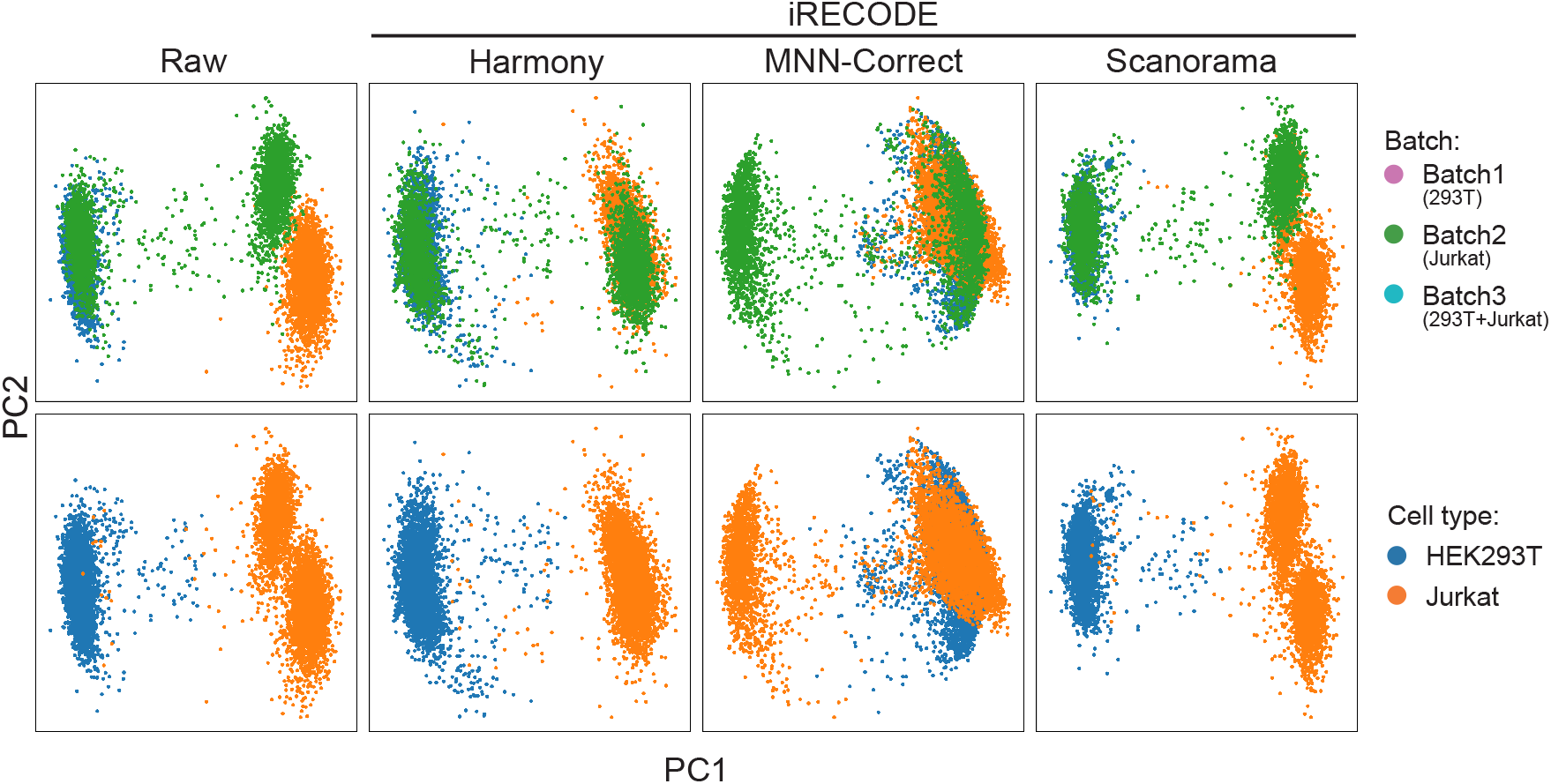
Compatibility of batch correction methodologies in iRECODE. This figure illustrates the performance of iRECODE incorporating three prominent batch correction algorithms―Harmony, MNN-Correct, and Scanorama―across datasets encompassing three batches and two cell lines (HEK293T and Jurkat). The first column represents the raw, uncorrected data, whereas the subsequent columns depict the data post-iRECODE using each method along the first two principal components (PC1 and PC2). iRECODE incorporating Harmony is notably effective in integrating the batches while maintaining cell type distinctions, as evidenced by the aligned clusters that segregate according to cell type rather than batch, underscoring its utility in harmonizing single-cell data while preserving biological variability.

**Figure S3:**
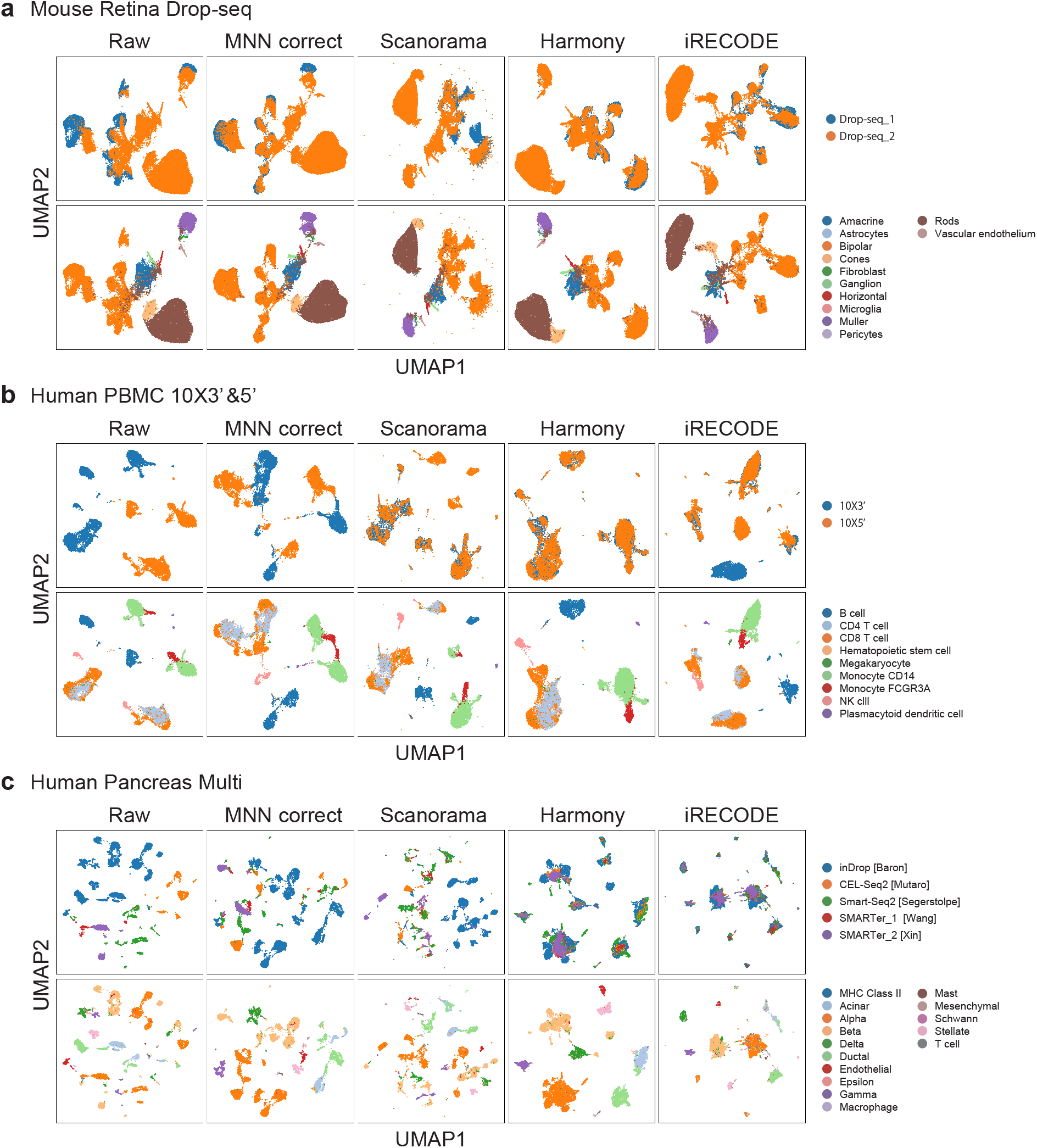
Comparative visualization of scRNA-seq data integration using iRECODE across multiple datasets and platforms. **a, b, c**, UMAP projections demonstrating batch and cell-type distribution within mouse retina data obtained via Drop-seq (**a**), human peripheral blood mononuclear cells (PBMC) sequenced using 10x Genomics’ 3’ and 5’ chemistry (**b**), and human pancreas cells sequenced using multiple single-cell technologies (**c**), with and without batch correction by various algorithms, including MNN correct, Scanorama, Harmony, and iRECODE.

**Figure S4:**
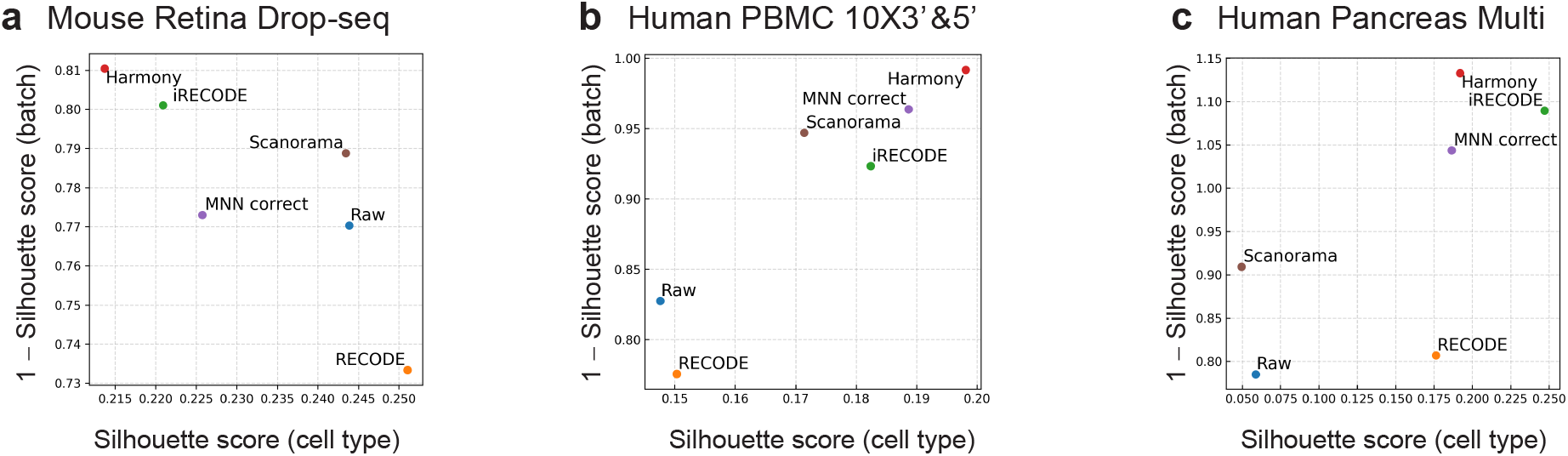
Quantitative assessment of batch correction efficacy using silhouette scores across multiple datasets in Fig S3. **a, b, c**, Scatter plot illustrating the silhouette scores for cell type versus batch correction in the mouse retina Drop-seq dataset (**a**), the human PBMC 10X 3’ and 5’ dataset (**b**), and the human pancreas multi-platform dataset (**c**). Various batch correction methods including MNN correct, Scanorama, Harmony, and iRECODE are compared against the raw and original RECODE. Each point represents the score of a particular method, with the proximity to the top right indicating superior batch correction performance. These evaluations provide insights into the comparative advantages of iRECODE in achieving coherent cell-type clustering while effectively mitigating batch variability.

**Figure S5:**
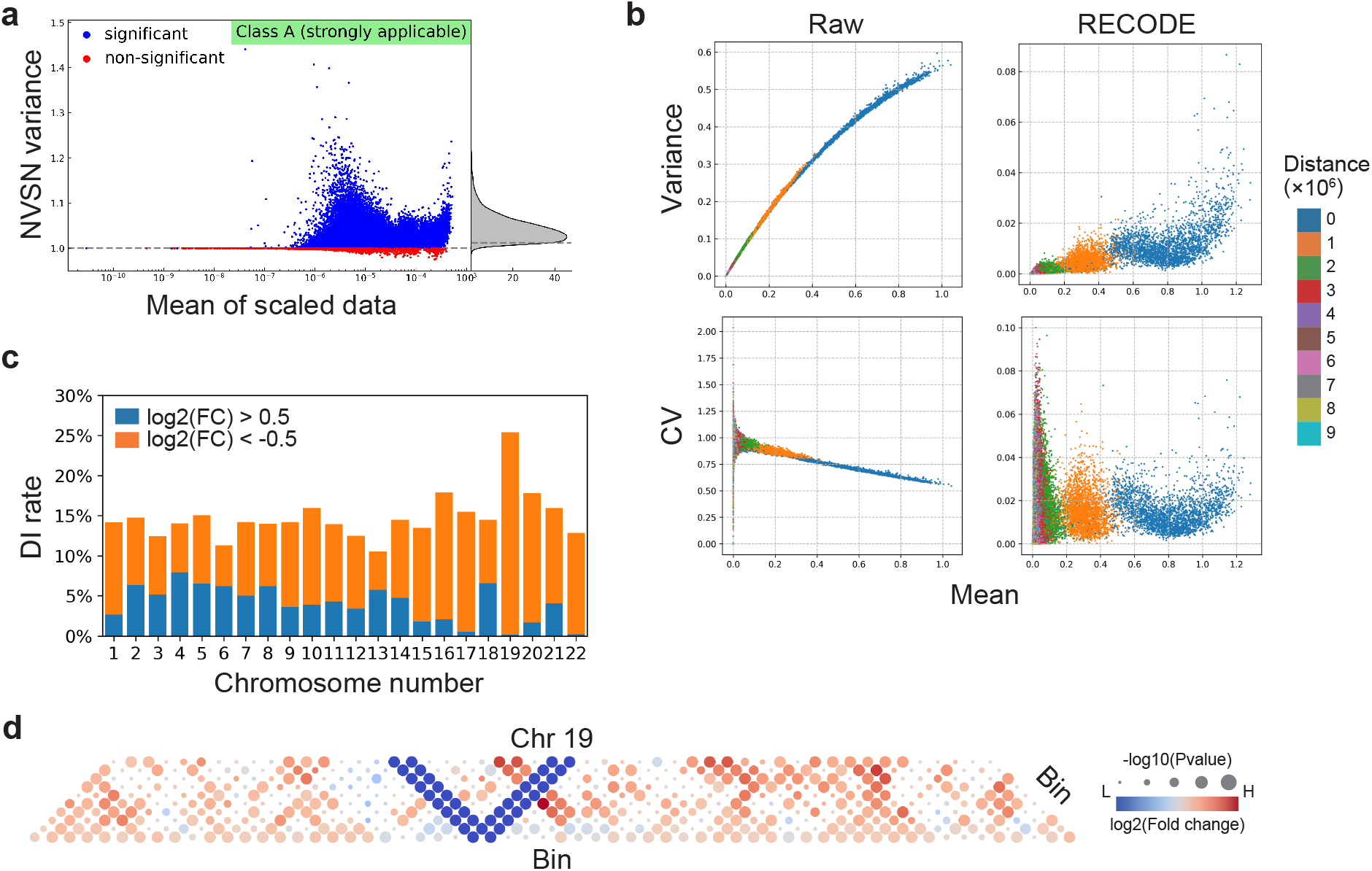
scHi-C data analysis via RECODE. **a**, NVSN plot demonstrating RECODE’s suitability for noise reduction of scHi-C data. **b**, Comparative plots of variance and coefficient of variation (CV) against the mean for raw and RECODE-processed scHi-C data, with color gradation representing inter-bin distances. **c**, Bar chart depicting the percentage of bins within differential interaction (DI) regions across all chromosomes, differentiated by the direction of log2 fold change (FC), providing insights into the chromosomal distribution of DIs defined in Fig 4e. **d**, Spatial representation of DIs along chromosome 19, with circle size indicating the magnitude of -log(*p*-value) and color denoting log2(FC), visualizing the regions of significant chromosomal interactions. These visualizations collectively quantify the impact of RECODE processing on the clarity and interpretability of scHi-C data, facilitating a deeper understanding of chromatin organization.

**Figure S6:**
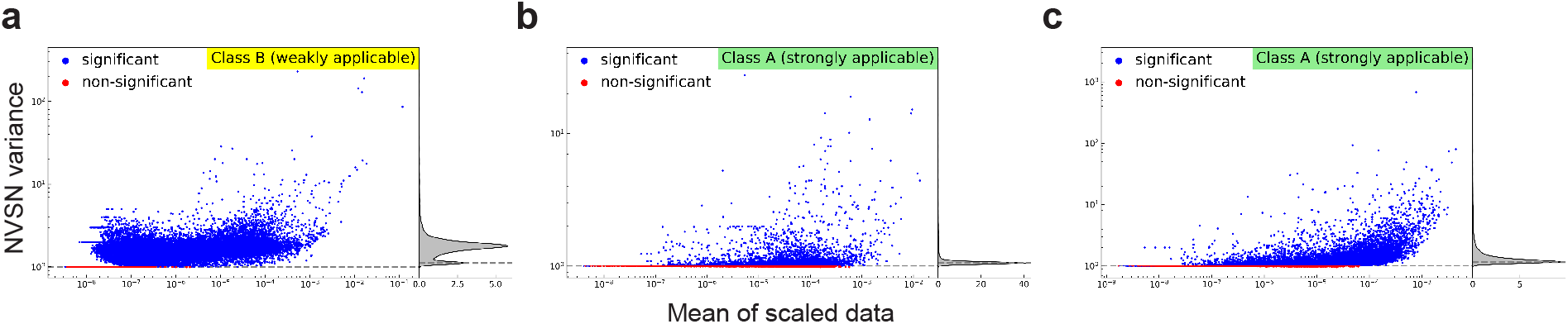
Assessment of the applicability of RECODE with spatial transcriptomics data. **a**, NVSN distribution for a mouse embryo spatial transcriptome dataset obtained via Stereo-seq, categorized as Class B (weakly applicable). **b**, NVSN distribution for a mouse brain dataset, and **c**, for a human colon cancer dataset, both acquired using the 10x Visium platform and classified as Class A (strongly applicable).

**Table S1:** Classification of the applicability of RECODE across diverse single-cell sequencing platforms.

|  |  | Strongly applicable | Weakly applicable | Inapplicable |
| --- | --- | --- | --- | --- |
| Transcriptome | 10x (RNA) | ✓ |  |  |
|  | Drop-seq | ✓ |  |  |
|  | Quartz-seq | ✓ |  |  |
|  | inDrop | ✓ |  |  |
|  | CEL-Seq2 | ✓ |  |  |
|  | DNBSEQ (MGISEQ) | ✓ |  |  |
|  | Smart-seq 2 |  | ✓ |  |
|  | Smart-seq 3 |  | ✓ |  |
|  | Fluidigm C1 |  | ✓ |  |
|  | SC3-seq |  | ✓ |  |
|  | TAS-Seq |  | ✓ |  |
|  | SMARTer |  |  | ✓ |
| Epigenome | 10x (ATAC) | ✓ |  |  |
|  | sci-Hi-C | ✓ |  |  |
| Multiome | 10x (RNA+ATAC) | ✓ |  |  |
|  | CITE-seq | ✓ |  |  |
| Spatial transcriptome | 10x Visium | ✓ |  |  |
|  | Stereo-seq |  | ✓ |  |

